## Supplementary Information for "Biogenesis of a bacterial metabolosome for propanediol utilization"

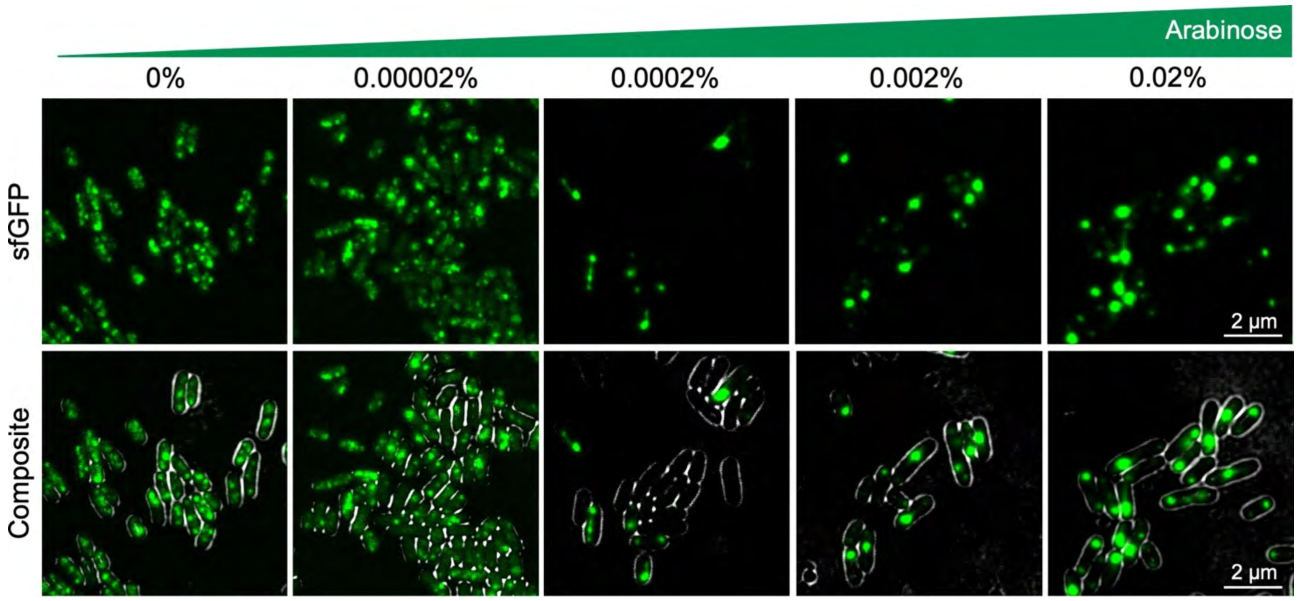


**Fig. S1. Fluorescence images show WT LT2 carrying pBAD-PduA-sfGFP grown in MIM+1,2-PD media at various arabinose concentrations (g·mL$^{-1}$).** Before fluorescence imaging, *Salmonella* cells were grown in a 2-mL Eppendorf tube shaken horizontally and aerobically at 37°C at 220 rpm until $OD_{600}$ reaching 1.0-1.2.

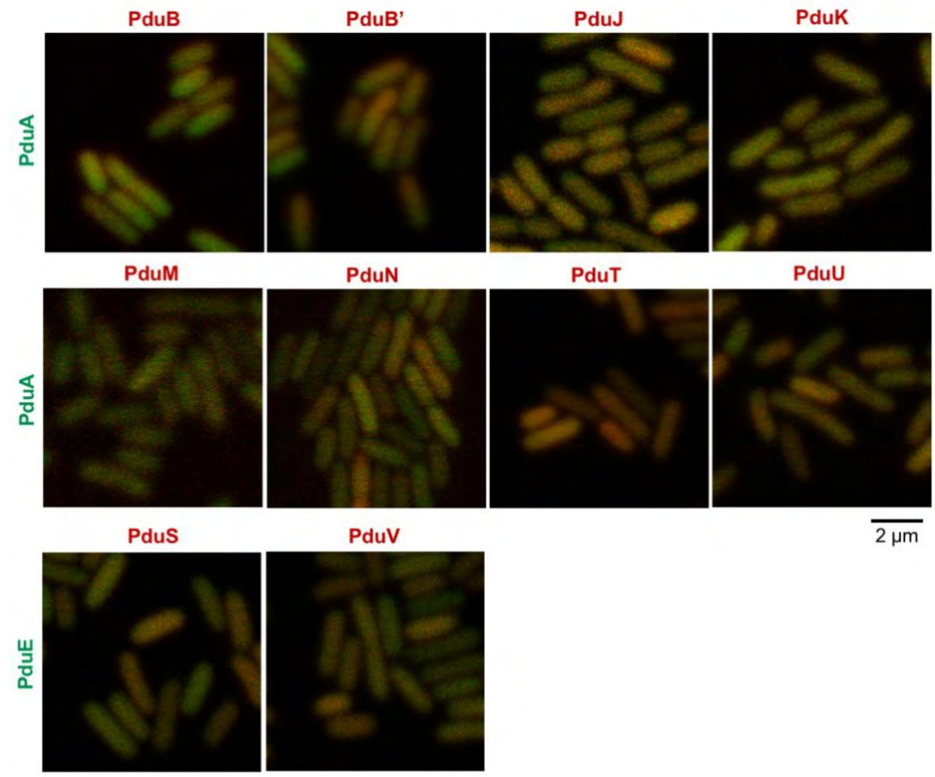


**Fig. S2. Pdu BMCs are not formed in the absence of 1,2-PD, as revealed by fluorescence images of the WT LT2 cells carrying pBAD (expressing fluorescently tagged PduA/E/B/B'/J/K/M/N/T/U/S/V proteins) grown in MIM-1,2-PD media.** Green represents the fluorescence of Pdu proteins tagged with sfGFP and red represents the fluorescence of proteins tagged with mCherry. Before fluorescence imaging, *Salmonella* cells were grown in a 2-mL Eppendorf tube shaken horizontally and aerobically at 37°C at 220 rpm until $OD_{600}$ reaching 1.0-1.2.

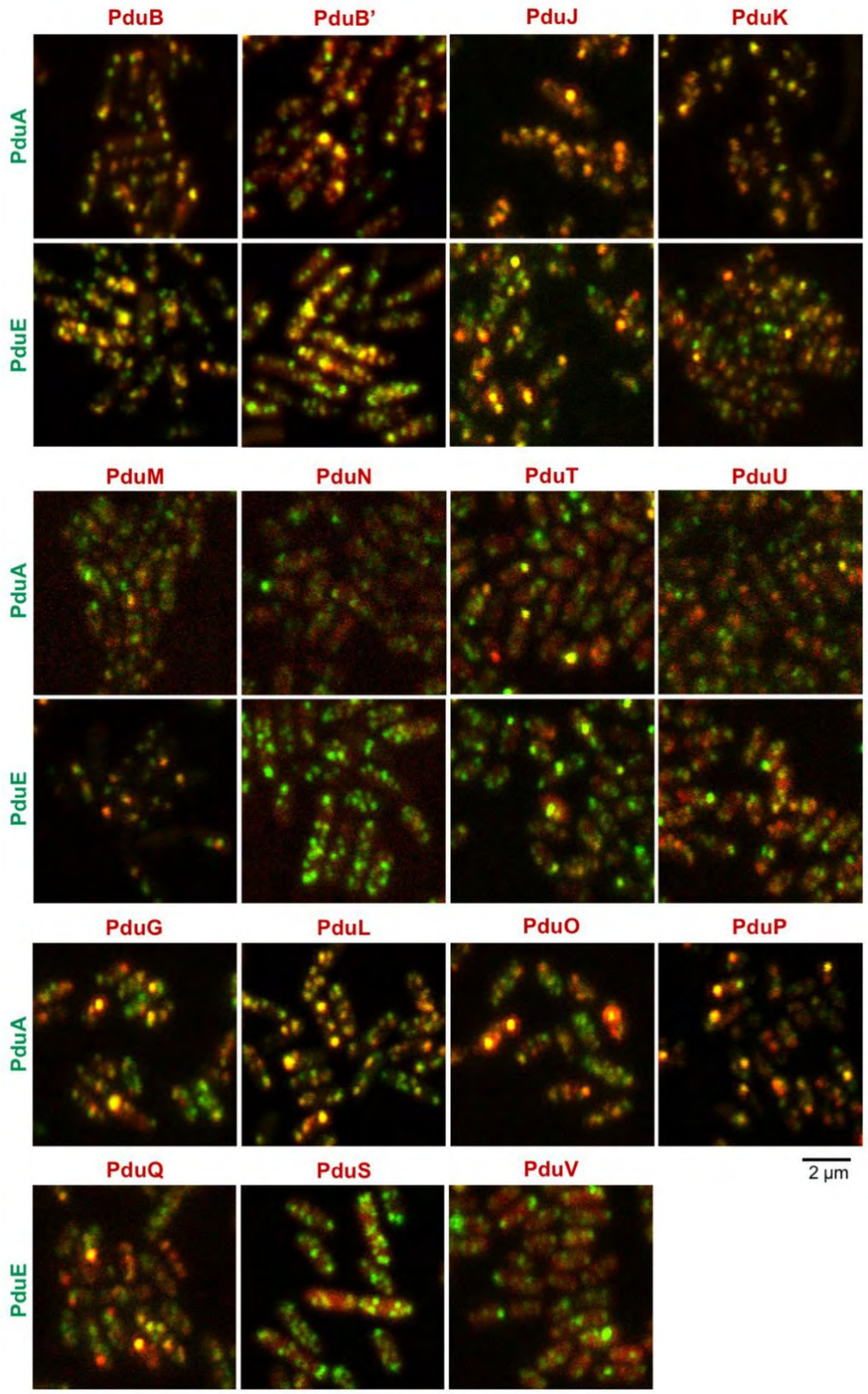


**Fig. S3. Pdu BMCs are formed in the presence of 1,2-PD.** Fluorescence images show WT LT2 carrying pBAD (expressing fluorescently tagged Pdu proteins) grown in MIM+1,2-PD media. Before fluorescence imaging, *Salmonella* cells were grown in a 2-mL Eppendorf tube shaken horizontally and aerobically at 37°C at 220 rpm until $OD_{600}$ reaching 1.0-1.2. Green represents the fluorescence of Pdu proteins tagged with sfGFP; red represents the fluorescence of proteins tagged with mCherry.

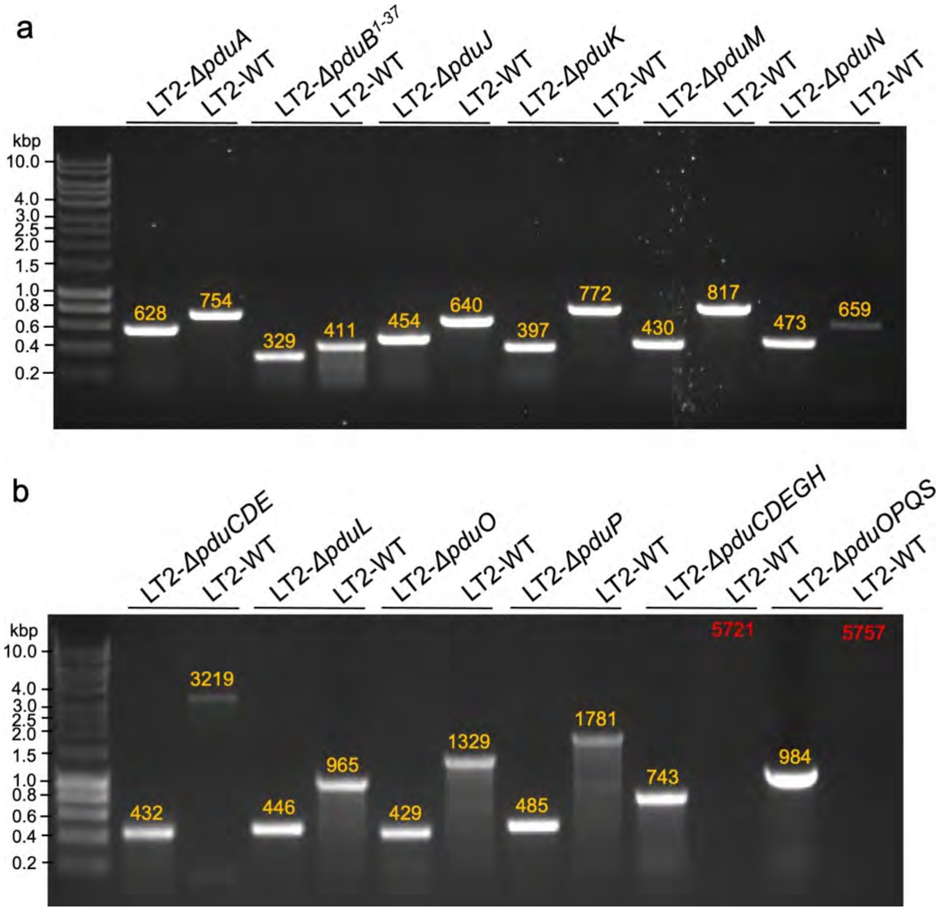


**Fig. S4. PCR-based confirmation of *S.* Typhimurium LT2 gene deletion mutants.** (**a**) and (**b**) correspond to shell and catalytic gene deletions, respectively. The sizes of the PCR products are indicated (bp, yellow). No band was detected in two WT strains due to the expected size (red) being too long to synthesize under these amplification conditions.

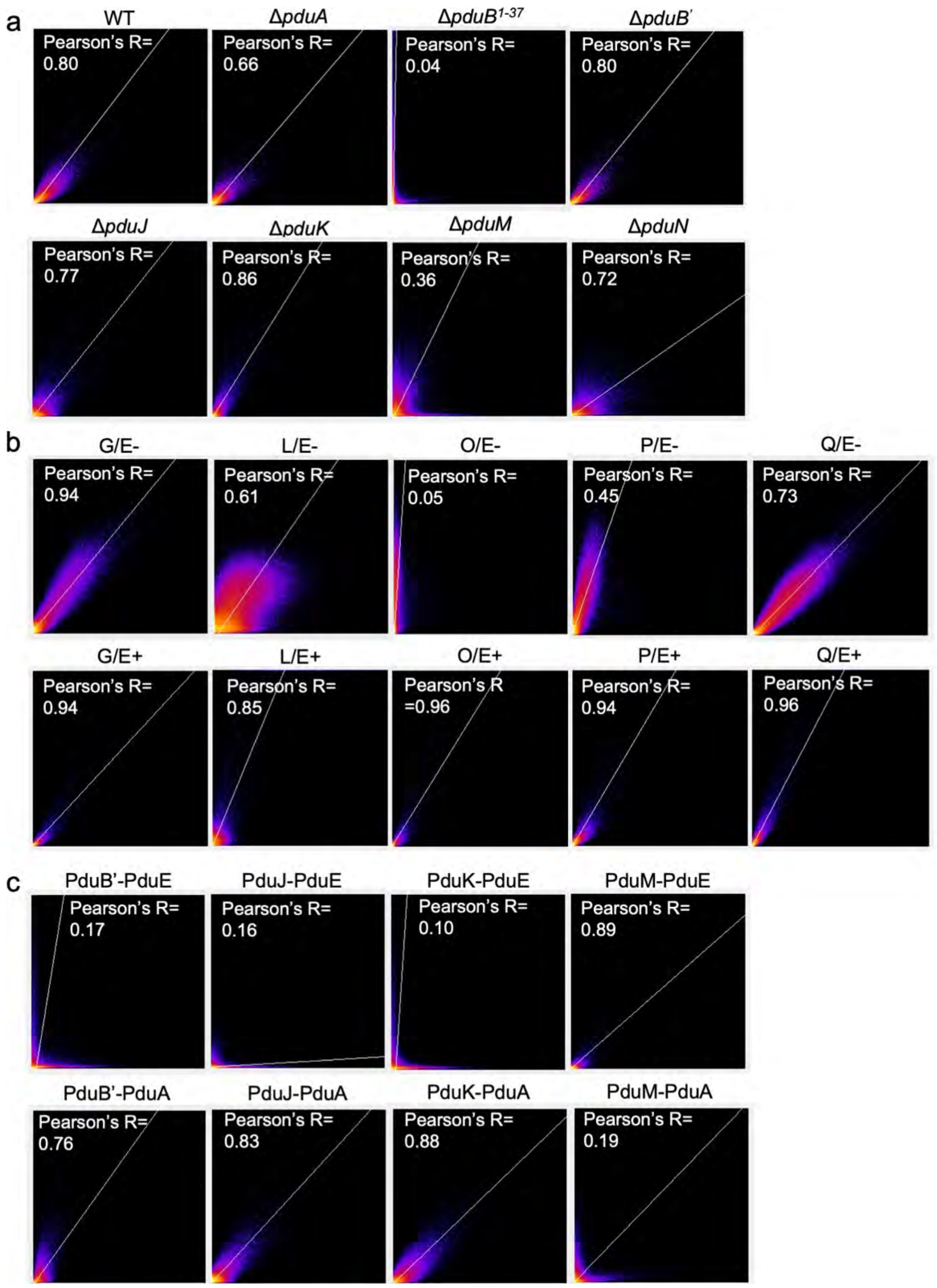


**Fig. S5. Scatterplot of colocalization analysis of data from Figs. 2 and 3.** (**a-c**) are representative scatterplots of Figs. 2b, 3b, and 3f, respectively. Scatterplots are generated by plotting the intensity value of each pixel of mCherry along the x-axis and the intensity value of the same pixel location of sfGFP on the y-axis using Coloc2 plugins in ImageJ. The scatterplots describe the relationship between the fluorescent signals. If the dots on the diagram appear as a cloud clustered on a line, a strong colocalization is indicated, and the Pearson's R is close to 1. If the scattered distribution of the pixels is close to both axes, a mutual exclusion is indicated, and the Pearson's R is near zero. Note: in panel (**b**), the first capital letter of the name is the name of the Pdu protein tagged with mCherry and the second capital letter of the name is the name of the Pdu protein tagged with sfGFP. '+' and '-' represents the presence and absence of 1,2-PD in the growth media, respectively. For example: 'G/E+' stands for PduG-mCherry/PduE-sfGFP in presence of 1,2-PD.

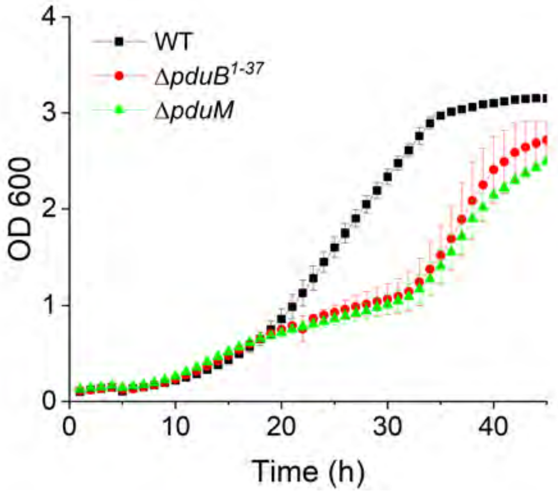


**Fig. S6. Growth curves of the *S.* Typhimurium LT2 WT and the Δ*pduB*$^{1-37}$ and Δ*pduM* mutants in the presence of 1,2-PD with saturating vitamin $B_{12}$ (150 nM).** The medium was NCE medium (containing 0.6% 1,2-PD; 0.3 mM each of leucine, isoleucine, threonine, and valine; 50 µM ferric citrate; 150 nM vitamin $B_{12}$). Growth curves were measured on a Growth Profiler 960 (EnzyScreen) under aerobic conditions.

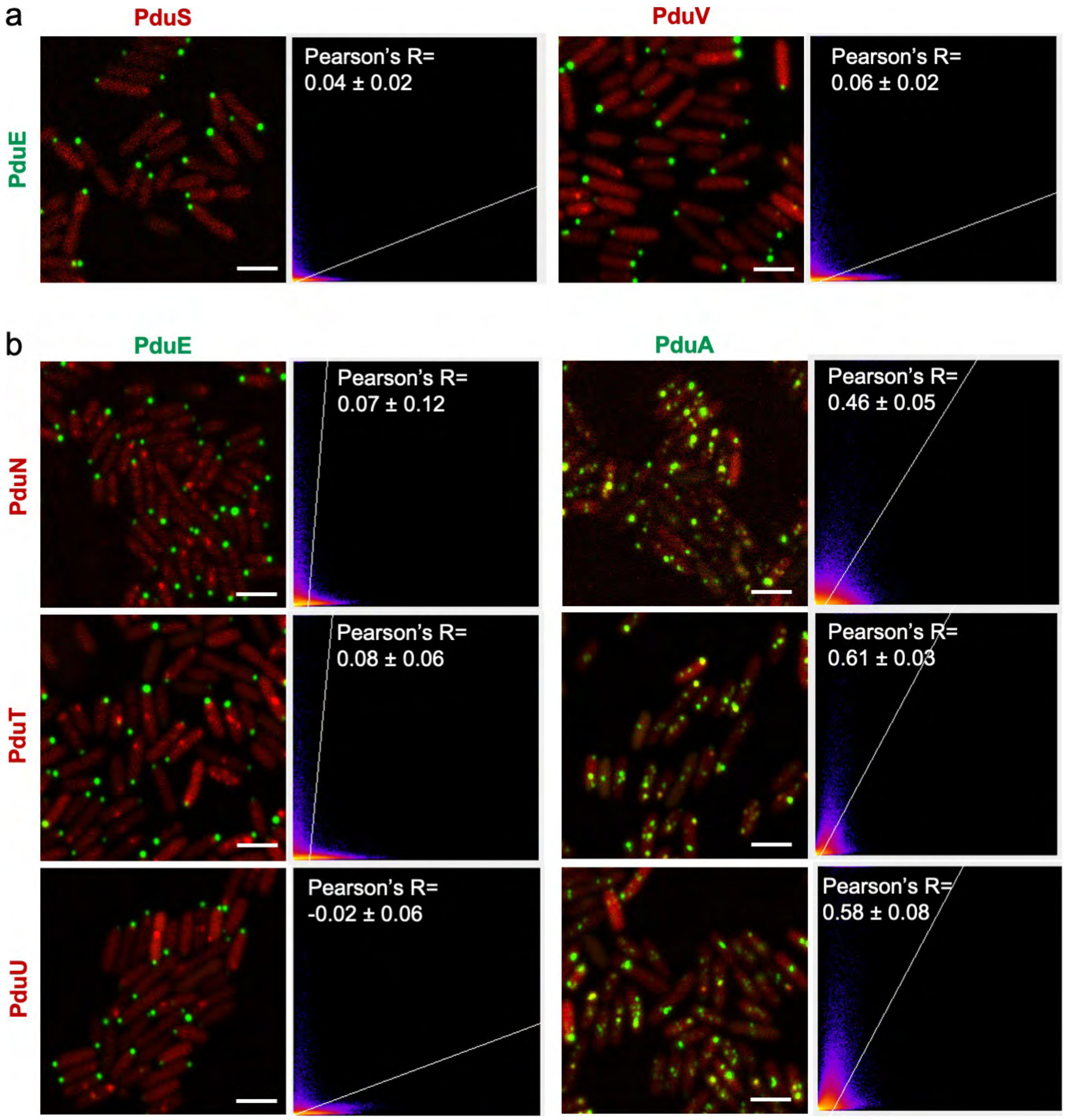


**Fig. S7. Minor Pdu enzymes (PduS/V) and minor Pdu shell proteins (PduN/T/U) are not colocalized with PduE (cargo).** (**a**) Fluorescence imaging of Δ*pduB*$^{1\text{-}37}$ expressing PduS-mCherry/PduE-sfGFP and PduV-mCherry/PduE-sfGFP (grown in MIM+1,2-PD media), and scatterplots of colocalization analysis. (**b**) Fluorescence imaging on Δ*pduB*$^{1\text{-}37}$ expressing minor shell protein (PduN, PduT and PduU) tagged with mCherry and PduE-sfGFP or PduA-sfGFP (grown in MIM+1,2-PD media), and scatterplots of colocalization analysis. Before fluorescence imaging, *Salmonella* cells were grown in a 2-mL Eppendorf tube shaken horizontally and aerobically at 37°C at 220 rpm until $OD_{600}$ reaching 1.0-1.2.

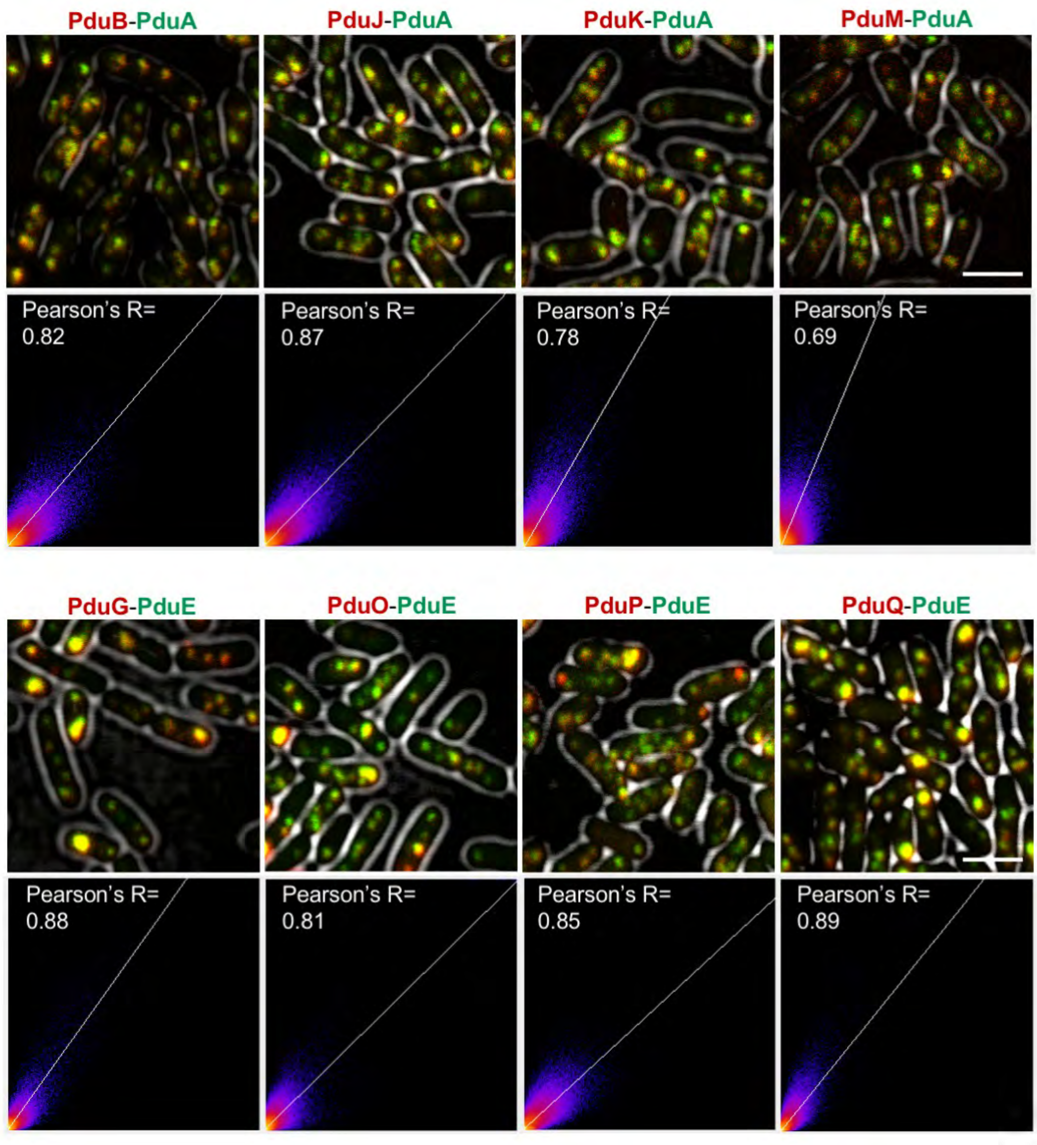


**Fig. S8. PduB' is not essential for assembly of the Pdu MCP.** Fluorescence imaging shows in presence of Pdu MCPs in Δ*pduB'* grown in MIM+1,2-PD media. Before fluorescence imaging, *Salmonella* cells were grown in a 2-mL Eppendorf tube shaken horizontally and aerobically at 37°C at 220 rpm until $OD_{600}$ reaching 1.0-1.2. Green represents the fluorescence of Pdu proteins tagged with sfGFP; red represents the fluorescence of proteins tagged with mCherry. Scatterplots are generated by plotting the intensity value of each pixel of mCherry along the x-axis and the intensity value of the same pixel location of sfGFP on the y-axis using Coloc2 plugins in ImageJ.

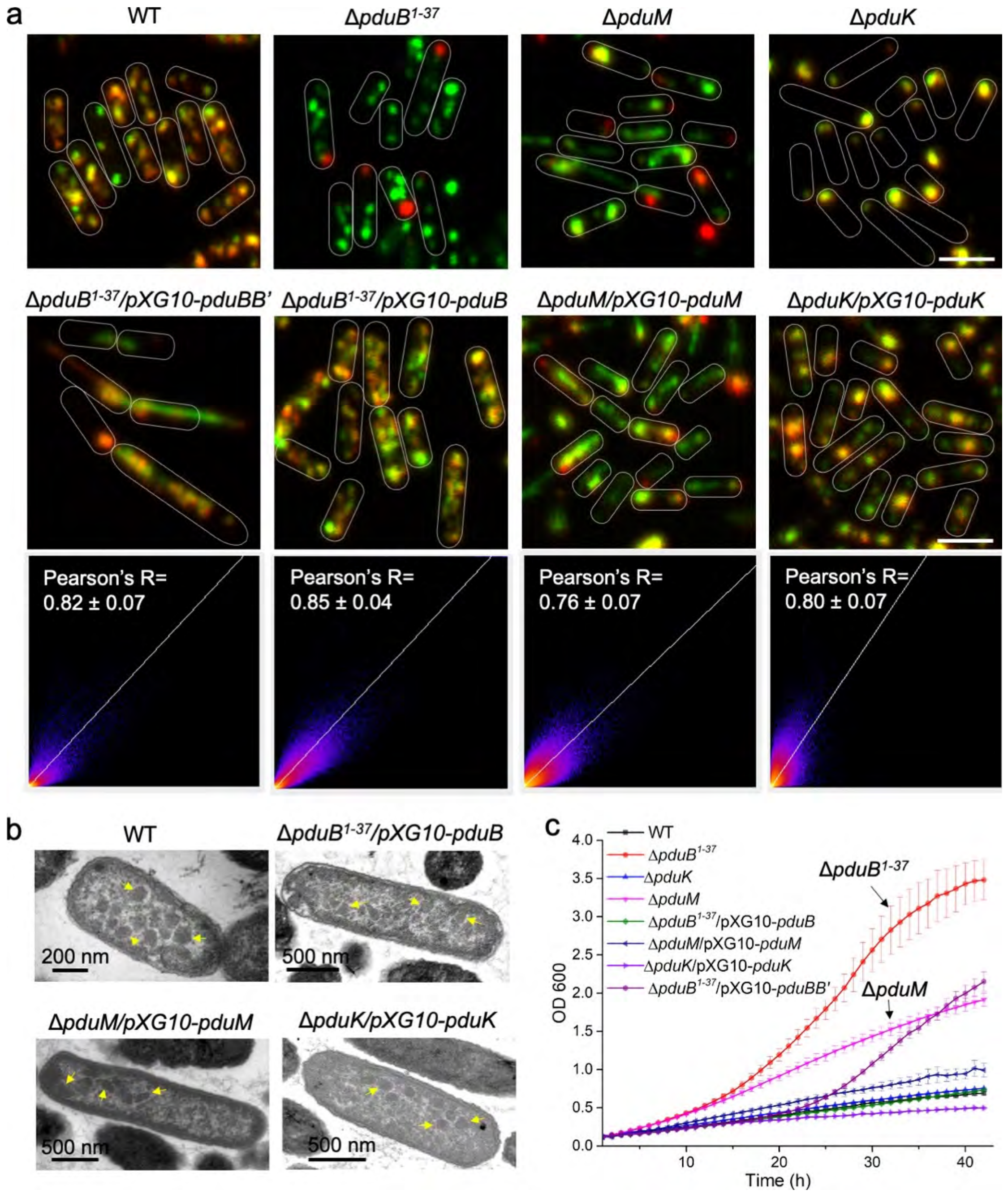


**Fig. S9. Successful complementation of PduB and PduK, and partial complementation of PduM.** (**a**) Fluorescence imaging on gene deletion mutants expressing deleted proteins from pXG10 plasmid grown in MIM+1,2-PD media, and scatterplots of colocalization analysis. Before fluorescence imaging, *Salmonella* cells were grown in a 2-mL Eppendorf tube shaken horizontally and aerobically at 37°C at 220 rpm until $OD_{600}$ reaching 1.0-1.2. (**b**) EM of WT and gene deletion mutants expressing deleted proteins from pXG10 plasmid grown in MIM+1,2-PD media. The Pdu MCP structures are indicated with yellow arrows. Before EM sample preparation, 10 mL *Salmonella* cells were grown aerobically in 50 mL Falcon tubes at 37°C at 220 rpm until $OD_{600}$ reaching 1.0-1.2. (**c**) Growth curves of gene deletion mutants carrying pXG10-based plasmids expressing the appropriate Pdu protein, during grown in 1,2-PD with limiting $B_{12}$ (20 nM). The medium was NCE medium (containing 0.6% 1,2-PD; 0.3 mM each of leucine, isoleucine, threonine, and valine; 50 µM ferric citrate; 20 nM vitamin $B_{12}$).

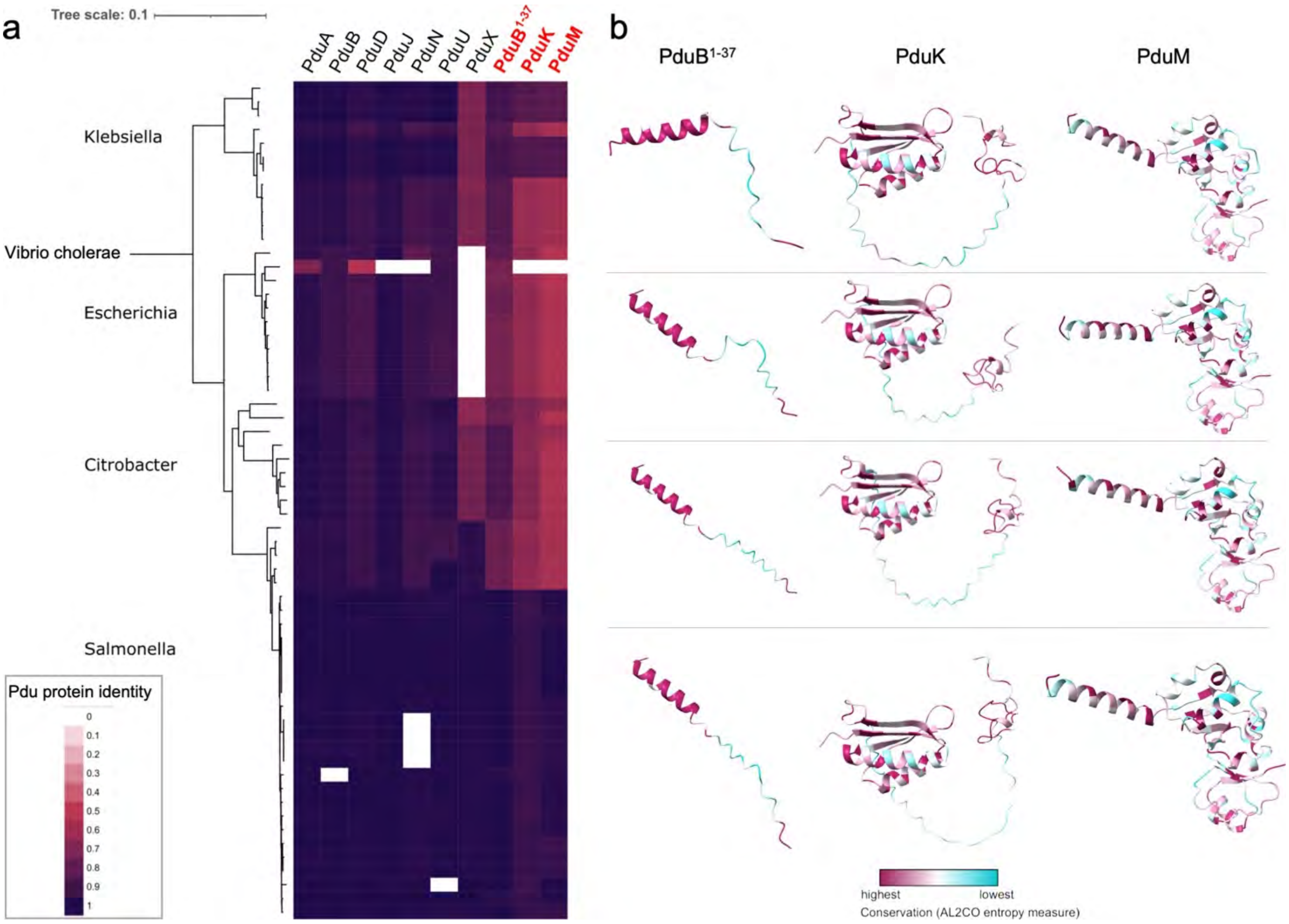


**Fig. S10. Similarities between the PduB$^{1\text{-}37}$, PduK and PduM proteins from four bacterial genera (*Klebsiella*, *Escherichia*, *Citrobacter,* and *Salmonella*) at the amino acid and structural levels.** (**a**) The phylogenetic tree was made from an alignment of 327 core genes from 61 bacterial genomes (see Supplementary File 1). The tree was rooted on the *Vibrio cholerae* MS6 genome as an outgroup. The heatmap shows the similarity of ten Pdu protein sequences of the 61 genomes comparing to *Salmonella* Typhimurium LT2. The levels of conservation of PduB$^{1\text{-}37}$, PduK, and PduM are mirrored by the conservation of other structural or essential Pdu proteins (PduA/B/D/J/N/U) encoded by the *pdu* operon. (**b**) The high structural similarity of proteins from different genera is inferred from AlphaFold 2 structural prediction. The structures are coloured by the conservation values of each residue in the alignment of 61 isolates. Four rows of the predicted structures are from four bacterial genera (from top to bottom, *Klebsiella*, *Escherichia*, *Citrobacter,* and *Salmonella*, respectively).

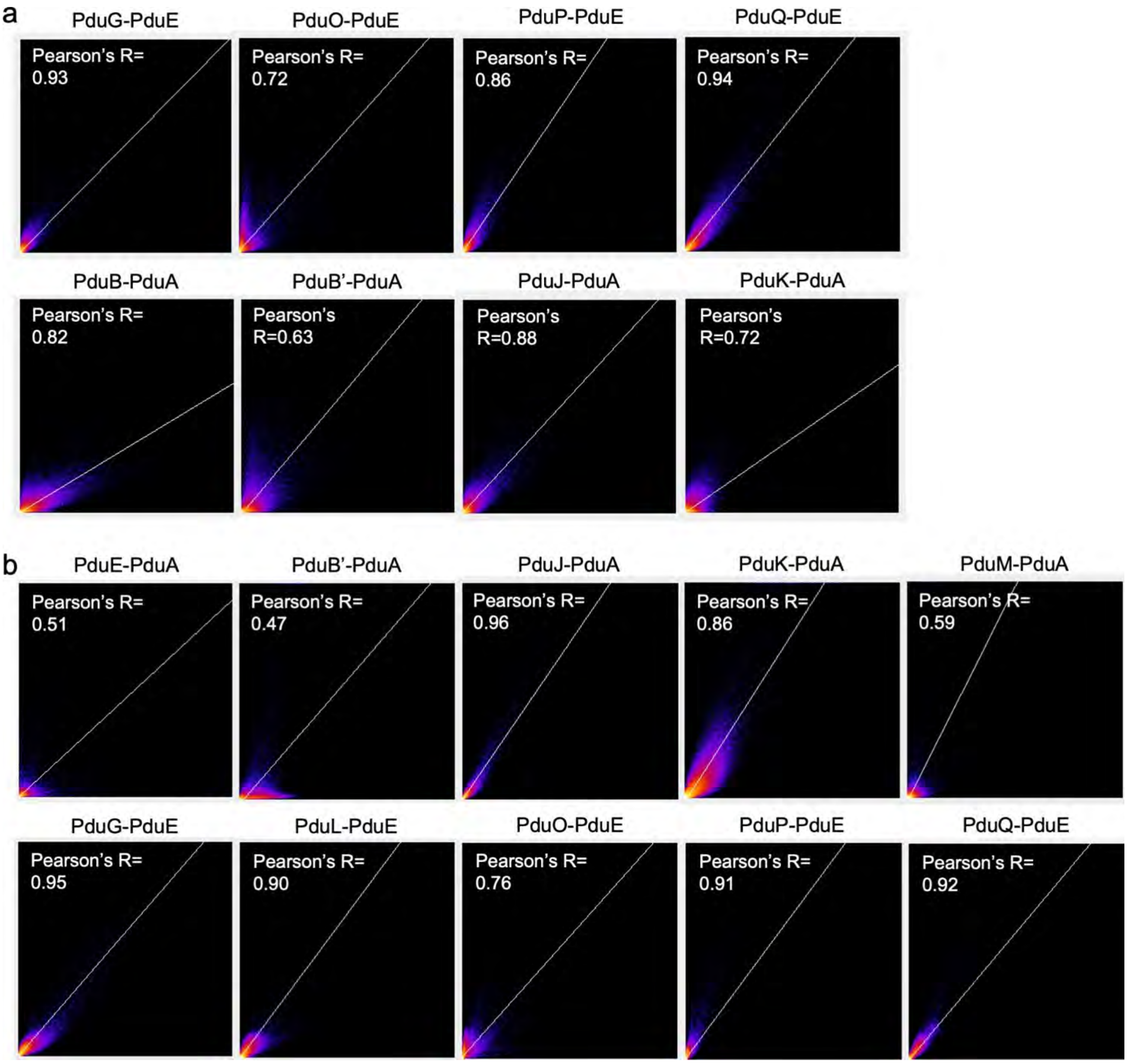


**Fig. S11. Scatterplot of colocalization analysis of data from Fig. 4.** (**a**) and (**b**) are representative scatterplots of Fig. 4B and 4F, respectively.

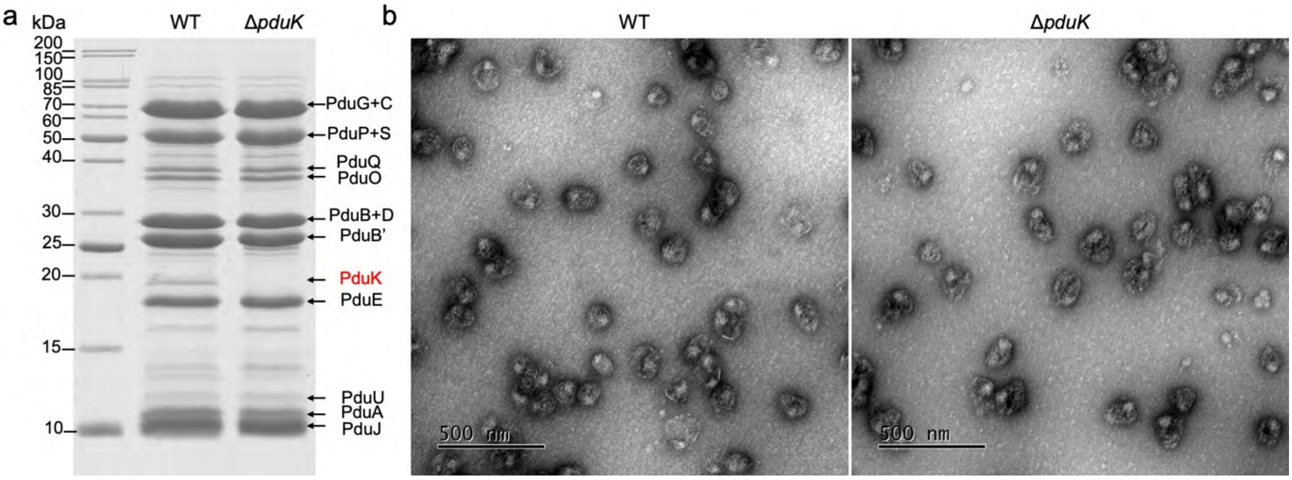


**Fig. S12. Isolation of Pdu BMCs from the *S*. Typhimurium LT2 WT and Δ*pduK* mutant.** (**a**) SDS-PAGE of isolated Pdu BMCs from the WT and Δ*pduK* cells. The Pdu BMCs were isolated from 400 mL cells grown aerobically in 1L flask in the MIM+1,2-PD medium ($OD_{600}$ = 1.0-1.2). (**b**) Negative-staining EM images of isolated Pdu MCPs from the WT and Δ*pduK* cells.

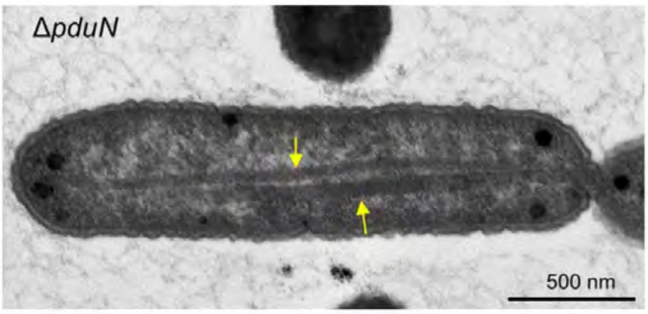


**Fig. S13. Thin-section EM of the Δ*pduN* strain revealing the elongated Pdu BMC structures (arrows) grown in the MIM+1,2-PD media.** Before EM sample preparation, 10 mL *Salmonella* cells were grown aerobically in 50 mL Falcon tubes at 37°C at 220 rpm until $OD_{600}$ reaching 1.0-1.2.

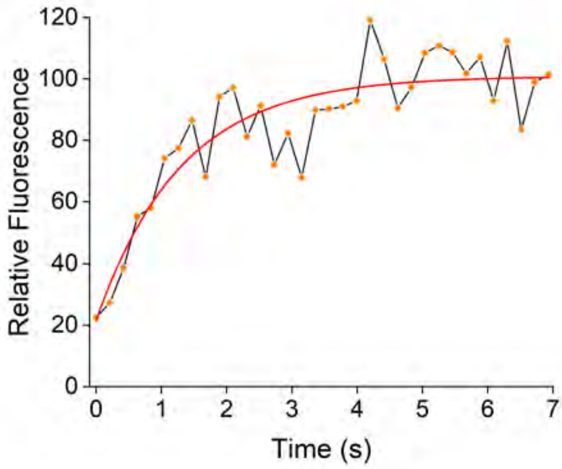


**Fig. S14. Time course of fluorescence recovery of bleached regions of free PduGH-sfGFP proteins.** The y-axis indicates fluorescence values relative to fluorescence intensity of the selected region prior to bleaching. The recovery of sfGFP fluorescence is shown as circles which were fitted to an exponential function.

**Table S1. Strains of *S.* Typhimurium LT2 derivatives and plasmids.** Relevant antibiotic resistances are indicated by $^R$: Ap, ampicillin; Km, kanamycin; Cm, chloramphenicol; Gm, gentamicin; Tc, tetracycline.

| Strains/Plasmids | Description | Origin |
|---|---|---|
| **LT2 derivatives:** | | |
| LT2 | LT2, WT | [1] |
| LT2-Δ*pduA* | Δ*pduA* | This study |
| LT2-*pduA*-sf*GFP* | LT2 derivative with PduA fused with sfGFP | This study |
| LT2-Δ*pduB*$^{1\text{-}37}$ | The N-terminus that differentiate PduB and PduB' was deleted | This study |
| LT2-Δ*pduB'* | The start codon for PduB' is replaced by GCT (alanine) | This study |
| LT2-Δ*pduJ* | Δ*pduJ* | This study |
| LT2-Δ*pduK* | Δ*pduK* | This study |
| LT2-Δ*pduM* | Δ*pduM* | This study |
| LT2-Δ*pduN* | Δ*pduN* | This study |
| LT2-Δ*pduCDE* | Δ*pduCDE* | This study |
| LT2-Δ*pduOPQS* | Δ*pduOPQS* | This study |
| LT2-Δ*pduB*$^{1\text{-}37}$/Δ*pduK* | Δ*pduB*$^{1\text{-}37}$ and Δ*pduK* double mutant | This study |
| LT2-Δ*pduB*$^{1\text{-}37}$/Δ*pduCDE* | Δ*pduB*$^{1\text{-}37}$ and Δ*pduCDE* double mutant | This study |
| LT2-Δ*pduB*$^{1\text{-}37}$/Δ*pduOPQS* | Δ*pduB*$^{1\text{-}37}$ and Δ*pduOPQS* double mutant | This study |
| ***E. coli* derivatives:** | | |
| *E. coli* S17-1 λ*pir* | *pro thi hsdR recA* chromosome::RP4-2 Tc::Mu Km::Tn7/λ*pir*; Tp$^R$, Sm$^R$ | [2] |
| **Plasmids:** | | |
| pKD4 | *aph*-cassette template plasmid; Km$^R$ | [3] |
| pKD3 | Template for amplification of Cm$^R$ cassette; Cm$^R$, Ap$^R$ | [3] |
| pCP20 | Plasmid carrying the Flp recombinase to remove kanamycin resistance from pKD13 derived resistance cassette insertions; Ap$^R$ | [3] |
| pSIM5-*tet* | λ Red recombination plasmid, temperature-inducible; Tc$^R$ | [4] |
| pEMG | Suicide plasmid; Km$^R$ | [5] |
| pEMG-pduB1-37_del | pEMG bearing a 1kb EcoRI-BamHI insert for deleting region pduB1-37; Km$^R$ | This study |
| pEMG-pduB M38A | pEMG bearing a 1kb EcoRI-BamHI insert for pduB M38A mutation; Km$^R$ | This study |
| pSW-2 | Plasmid for m-toluate–inducible expression of the I-SceI enzyme; Gm$^R$ | [5] |
| pBAD/*Myc*-His | Vector for dose-dependent expression of recombinant proteins; Ap$^R$ | Invitrogen |

| | | |
|---|---|---|
| pBAD-*EA* | *pduE::mCherry-pduA::sfGFP* cloned into pBAD/*Myc*-His at NcoI and HindIII sites; $Ap^R$ | This study |
| pBAD-*BA* | *pduB::mCherry-pduA::sfGFP* cloned into pBAD/*Myc*-His at NcoI and HindIII sites; $Ap^R$ | This study |
| pBAD-*B'A* | *pduB'::mCherry-pduA::sfGFP* cloned into pBAD/*Myc*-His at NcoI and HindIII sites; $Ap^R$ | This study |
| pBAD-*JA* | *pduJ::mCherry-pduA::sfGFP* cloned into pBAD/*Myc*-His at NcoI and HindIII sites; $Ap^R$ | This study |
| pBAD-*KA* | *pduK::mCherry-pduA::sfGFP* cloned into pBAD/*Myc*-His at NcoI and HindIII sites; $Ap^R$ | This study |
| pBAD-*MA* | *pduM::mCherry-pduA::sfGFP* cloned into pBAD/*Myc*-His at NcoI and HindIII sites; $Ap^R$ | This study |
| pBAD-*NA* | *pduN::mCherry-pduA::sfGFP* cloned into pBAD/*Myc*-His at NcoI and HindIII sites; $Ap^R$ | This study |
| pBAD-*TA* | *pduT::mCherry-pduA::sfGFP* cloned into pBAD/*Myc*-His at NcoI and HindIII sites; $Ap^R$ | This study |
| pBAD-*UA* | *pduU::mCherry-pduA::sfGFP* cloned into pBAD/*Myc*-His at NcoI and HindIII sites; $Ap^R$ | This study |
| pBAD-*BE* | *pduB::mCherry-pduE::sfGFP* cloned into pBAD/*Myc*-His at NcoI and HindIII sites; $Ap^R$ | This study |
| pBAD-*B'E* | *PduB'::mCherry-pduE::sfGFP* cloned into pBAD/*Myc*-His at NcoI and HindIII sites; $Ap^R$ | This study |
| pBAD-*JE* | *pduJ::mCherry-pduE::sfGFP* cloned into pBAD/*Myc*-His at NcoI and HindIII sites; $Ap^R$ | This study |
| pBAD-*KE* | *pduK::mCherry-pduE::sfGFP* cloned into pBAD/*Myc*-His at NcoI and HindIII sites; $Ap^R$ | This study |
| pBAD-*ME* | *pduM::mCherry-pduE::sfGFP* cloned into pBAD/*Myc*-His at NcoI and HindIII sites; $Ap^R$ | This study |
| pBAD-*NE* | *pduN::mCherry-pduE::sfGFP* cloned into pBAD/*Myc*-His at NcoI and HindIII sites; $Ap^R$ | This study |
| pBAD-*TE* | *pduT::mCherry-pduE::sfGFP* cloned into pBAD/*Myc*-His at NcoI and HindIII sites; $Ap^R$ | This study |
| pBAD-*UE* | *pduU::mCherry-pduE::sfGFP* cloned into pBAD/*Myc*-His at NcoI and HindIII sites; $Ap^R$ | This study |
| pBAD-*GE* | *pduG::mCherry-pduE::sfGFP* cloned into pBAD/*Myc*-His at NcoI and HindIII sites; $Ap^R$ | This study |
| pBAD-*LE* | *pduL::mCherry-pduE::sfGFP* cloned into pBAD/*Myc*-His at NcoI and HindIII sites; $Ap^R$ | This study |
| pBAD-*OE* | *pduO::mCherry-pduE::sfGFP* cloned into pBAD/*Myc*-His at NcoI and HindIII sites; $Ap^R$ | This study |
| pBAD-*PE* | *pduP::mCherry-pduE::sfGFP* cloned into pBAD/*Myc*-His at NcoI and HindIII sites; $Ap^R$ | This study |
| pBAD-*QE* | *pduQ::mCherry-pduE::sfGFP* cloned into pBAD/*Myc*-His at NcoI and HindIII sites; $Ap^R$ | This study |
| pBAD-*SE* | *pduS::mCherry-pduE::sfGFP* cloned into pBAD/*Myc*-His at NcoI and HindIII sites; $Ap^R$ | This study |

| | | |
|---|---|---|
| pBAD-*VE* | *pduV::mCherry-pduE::sfGFP* cloned into pBAD/*Myc*-His at NcoI and HindIII sites; $Ap^R$ | This study |
| pBAD-*pduCDE-sfGFP* | *pduCDE::sfGFP* cloned into pBAD/*Myc*-His at NcoI and HindIII sites; $Ap^R$ | This study |
| pBAD-*pduGH-sfGFP* | *pduGH::sfGFP* cloned into pBAD/*Myc*-His at NcoI and HindIII sites; $Ap^R$ | This study |
| pBAD-*pduE-sfGFP* | *pduE::sfGFP* cloned into pBAD/*Myc*-His at NcoI and HindIII sites; $Ap^R$ | This study |
| pBAD-*pduA-sfGFP* | *pduA::sfGFP* cloned into pBAD/*Myc*-His at NcoI and HindIII sites; $Ap^R$ | This study |
| pXG10-SF | Plasmid served as the backbone for complementation experiments; pSC101* origin of replication; $P_{LtetO\text{-}1}$ promoter; $Cm^R$ | [6] |
| pXG10-*pduB* | Plasmid for expression of PduB (M38A); $Cm^R$ | This study |
| pXG10-*pduBB'* | Plasmid for expression of PduBB'; $Cm^R$ | This study |
| pXG10-*pduM* | Plasmid for expression of PduM; $Cm^R$ | This study |
| pXG10-*pduK* | Plasmid for expression of PduK; $Cm^R$ | This study |

**Table S2. ssDNA Oligonucleotides used in this study.**

| Primers | Sequence (5'→3') |
|---|---|
| pduA_del_F | TCTTATAGTCCCAACTATCGGAACACTCCATGCGAGGTCTTTATGGTGT AGGCTGGAGCTGCTTC |
| pduA_del_R | GTTCCACCAGCTCATTGCTGCTCATTGGCTAATTCCCTTCGGTAACATAT GAATATCCTCCTTAG |
| pduA_up | AAATATTGCACAAGCCAACTTATC |
| pduA_down | GGCCCAGGGTATCGCCAATG |
| pduA_sfGFP_F | CCTCACACCGATGTAGAAAAAATCTTACCGAAGGGAATTAGCCAAGGA TCCGCTGGCTCCGCTGC |
| pduA_sfGFP_R | TCTGTTCCACCAGCTCATTGCTGCTCATTGGCTAATTCCCTTCGGGTGTA GGCTGGAGCTGCTTC |
| pduB1-37_del_F1 | AGGGATAACAGGGTAATCTGAATTGCACAAGCCAACTTATCAATTTCGA |
| pduB1-37_del_R1 | CCGTCTCTCGTATAGGTTGTCAGCTCATTGGCTAATTCCC |
| pduB1-37_del_F2 | GGGAATTAGCCAATGAGCTGACAACCTATACGAGAGACGG |
| pduB1-37_del_R2 | CCTGCAGGTCGACTCTAGAGGATCACTTCAGCGCGGTATCGGCC |
| pduB_up | TACACGTCATCCCACGCCCT |
| pduB_down | GTATCGCGCGGCAGCTCAAT |
| pduB M38A-F1 | AGGGATAACAGGGTAATCTGAATTCTGATGCTCAACAGCAAGTC |
| pduB M38A-R1 | AAACTGCAGCTTTTTCTGCAGCAGCCGTCTCTCGTATAG |
| pduB M38A-F2 | CTATACGAGAGACGGCTGCTGCAGAAAAAGCTGCAGTTT |
| pduB M38A-R2 | CCTGCAGGTCGACTCTAGAGGATCACTTCAGCGCGGTATCGGCC |
| pduJ_del_F | CCCTTTCGGGATCTCCATGCTTAATCACAGGAGAACGGCAGTATGGTGT AGGCTGGAGCTGCTTC |
| pduJ_del_R | GCGGTGCTCCTTATTCGCCATCGATTAGGCTGATTTCGGCATATGAATAT CCTCCTTAG |

| | |
|---|---|
| pduJ_up | GATCGACACTCGCTGGTCGT |
| pduJ_down | CCTGAACGGAGGCCACATCA |
| pduK_del_F | GATGTTGAGGCCATTTTACCGAAATCAGCCTAATCGATGGTGTAGGCTG GAGCTGCTTC |
| pduK_del_R | GCAGAAGCTCTTTATCCATTACGCTTCACCTCGCTTGCCCATATGAATAT CCTCCTTAG |
| pduK_up | TCATGGTTCGCGGCGATGTC |
| pduK_down | GATGGGATGGCCGGGAAACA |
| pduM_del_F | CCCGCATGCCTTTGCCCGGCTGGTAGGCCCGCGATGAACGTGTAGGCTG GAGCTGCTTC |
| pduM_del_R | CGTGACTCGTGCCAGATGCATGATTTACTCCTGCTTAATCATATGAATAT CCTCCTTAG |
| pduM_up | GCGGGCTGATTTTCAACAAC |
| pduM_down | GCCGCTGAGCAAAACCAGTT |
| pduN_del_F | GCCAATGCGCGGAATATTCAATTAATTAAGCAGGAGTAAATCATGGTGT AGGCTGGAGCTGCTTC |
| pduN_del_R | GCCAGCGTCACCTGTTCGGGTATAAATCGCCATAACCGCCCCTTACATA TGAATATCCTCCTTAG |
| pduN_up | TGCCGCTGGTATTCACCGAT |
| pduN_down | CTGCTGGATGGCCTCGAGTA |
| pduCDE_del_F | CGTCCGTCCTACATCTGATACCCACGAGGCTGATTCATGGTGTAGGCTG GAGCTGCTTC |
| pduCDE_del_R | GCCAGCTATATATCGCATACGAAATCCTTAATCGTCGCCCATATGAATA TCCTCCTTAG |
| pduCDE_up | CGGCACCAGCTTTAGTAACG |
| pduCDE_down | TCCTGAATGCCGAACACGTT |
| pduCDEGH_del_R | TCCCAGTGCGTTATTCATACTGCCGTTCTCCTGTGATTACATATGAATAT CCTCCTTAG |
| pduL_del_F | GCTTTGCATTCATTCCGGCAAGCGAGGTGAAGCGTAATGCATATGAATA TCCTCCTTAG |
| pduL_del_R | GCAGGGTTTCGCCGTTCATCGCGGGCCTACCAGCCGGGCGTGTAGGCTG GAGCTGCTTC |
| pduL_up | CCTGAGCCTGAAGCGTCAG |
| pduL_down | GCAGGTCGATGAGCAGGAT |
| pduO_del_F | GGCATTGTAGATACGCTTTCGTGTTAAGGGGCGGTTATGGTGTAGGCTG GAGCTGCTTC |
| pduO_del_R | CGAGTTCAGAAGTATTCATTGATGAGTTCCCACGTTAATCATATGAATA TCCTCCTTAG |
| pduO_up | ATGAAGTGGCCGTGGACT |
| pduO_down | AGCGGGCACTGCTGATAAC |
| pduP_del_F | CGCCATCGCGGCTATTAACGTGGGAACTCATCAATGAATGTGTAGGCTG GAGCTGCTTC |

| | |
|---|---|
| pduP_del_R | GTAGTGAGAAGGTATTCATCGCGACCTCAGTTAGCGAATCATATGAATATCCTCCTTAG |
| pduP_up | GCTGAGCGATGTCGTTCA |
| pduP_down | GACGCTGATGCGGTTATCTG |
| pduOPQS_del_R | TTCCTATAGCCTGAGACATGGTTAACCTCTTACAACAGTCATATGAATATCCTCCTTAG |
| pduOPQS_down | TGCTTCGCTCGGCGTGATGG |
| pXG10-F | TCTAGAGGCATCAAATAAAACGAAAG |
| pXG10-R | ATGCATGTGCTCAGTATCTCTATCAC |
| pXG10-pduB-F | GTGATAGAGATACTGAGCACATGCATTGTAGAAAAAATCTTACCGAAGGGAATTAGCC |
| pXG10-pduB-R | CTTTCGTTTTATTTGATGCCTCTAGATCAGATGTAGGACGGACGATCG |
| pXG10-pduM-F | GTGATAGAGATACTGAGCACATGCATCCCGCATGCCTTTGCCCG |
| pXG10-pduM-R | CTTTCGTTTTATTTGATGCCTCTAGATTACTCCTGCTTAATTAATTGAATATTCCG |
| pXG10-pduK-F | GTGATAGAGATACTGAGCACATGCATGTTGAGGCCATTTTACCGAAAT |
| pXG10-pduK-R | CTTTCGTTTTATTTGATGCCTCTAGATTACGCTTCACCTCGCTTGC |
| pBAD-CDE-sfGFP-F | GGGCTAACAGGAGGAATTAACCATGAGATCGAAAAGATTTGAAGCAC |
| pBAD-CDE-sfGFP-R | GAGATGAGTTTTTGTTCTACGTATTATTTGTAGAGCTCATCCATGCC |
| pBAD-GH-sfGFP-F1 | GGGCTAACAGGAGGAATTAACCATGCGATATATAGCTGGCATTGACA |
| pBAD-GH-sfGFP-R1 | GGAGCCAGCGGATCCAGCATGGAGATCCCGA |
| pBAD-GH-sfGFP-F2 | ATGCTGGATCCGCTGGCTCCG |
| pBAD-GH-sfGFP-R2 | GAGATGAGTTTTTGTTCTACGTAAGCTT |
| pBAD-EA-F1 | GGGCTAACAGGAGGAATTAACCATGAATACCGACGCAATTGAATCG |
| pBAD-EA-R1 | AGAACCAGCAGCGGAGCCAGCGGATCCATCGTCGCCTTTGAGTTTTTA |
| pBAD-EA-F2 | CTGGCTCCGCTGCTGGTTCTGGCGAATTCGTGAGCAAGGGCGAGGAG |
| pBAD-EA-R2 | CTCCTGTTAGCCCCTACTTGTACAGCTCGTCCATGCCGCC |
| pBAD-EA-F3 | CAAGTAGGGGCTAACAGGAGGAATTAACCATGCAACAAGAAGCACTAGG |
| pBAD-EA-R3 | GAGATGAGTTTTTGTTCTACGTATTATTTGTAGAGCTCATCCATGCC |
| pBAD-PA-F1 | GGGCTAACAGGAGGAATTAACCATGAATACTTCTGAACTCGAAACCCTGATTCGCA |
| pBAD-PA-R1 | CTTGCTCACGAATTCGCCAGAACCAGCAGCGGAGCCAGCGGATCCGCGAATAGAAAAGCC |
| pBAD-PA-F2 | CTGGCGAATTCGTGAGCAAG |
| pBAD-PA-R2 | CTGAGATGAGTTTTTGTTCTACGTA |

| | |
|---|---|
| pBAD-PE-F1 | GGGCTAACAGGAGGAATTAACCATGAATACTTCTGAACTCGAAACCCTGATTCGCA |
| pBAD-PE-R1 | CTTGCTCACGAATTCGCCAGAACCAGCAGCGGAGCCAGCGGATCCGCGAATAGAAAAGCC |
| pBAD-PE-F2 | CTGGCGAATTCGTGAGCAAGGGCGA |
| pBAD-PE-R2 | TTGCGTCGGTATTCATGGTTAATTCCTCCTGTTAGCCCC |
| pBAD-PE-F3 | AACCATGAATACCGACGCAATTGAATCGATGGTCC |
| pBAD-PE-R3 | GAGATGAGTTTTTGTTCTACGTATTATTTGTAGAGCTCATCCA |
| pBAD-J-F1 | GGGCTAACAGGAGGAATTAACCATGAATAACGCACTGGGACTGGTTG |
| pBAD-J-R1 | CTTGCTCACGAATTCGCCAGAACCAGCAGCGGAGCCAGCGGATCCGGCTGATTTCGGTAA |
| pBAD-GE-F1 | GGGCTAACAGGAGGAATTAACCATGCGATATATAGCTGGCATTGACATCGGTAAC |
| pBAD-GE-R1 | CTTGCTCACGAATTCGCCAGAACCAGCAGCGGAGCCAGCGGATCCCTGTCCATGCGCAAA |
| pBAD-LE-F1 | GGGCTAACAGGAGGAATTAACCATGGATAAAGAGCTTCTGCAATCAACGGTC |
| pBAD-LE-R1 | CTTGCTCACGAATTCGCCAGAACCAGCAGCGGAGCCAGCGGATCCTCGCGGGCCTACCAG |
| pBAD-OE-F1 | GGGCTAACAGGAGGAATTAACCATGGCGATTTATACCCGAACAGGTGACG |
| pBAD-OE-R1 | CTTGCTCACGAATTCGCCAGAACCAGCAGCGGAGCCAGCGGATCCTTGATGAGTTCCCAC |
| pBAD-QE-F1 | GGGCTAACAGGAGGAATTAACCATGAATACCTTCTCACTACAAACGCGGTTGTA |
| pBAD-QE-R1 | CTTGCTCACGAATTCGCCAGAACCAGCAGCGGAGCCAGCGGATCCTAGCAGTTCCTCCAG |
| pBAD-SE-F1 | GGGCTAACAGGAGGAATTAACCATGAGCACCGCCATCAACAGCGTT |
| pBAD-SE-R1 | CTTGCTCACGAATTCGCCAGAACCAGCAGCGGAGCCAGCGGATCCACCTCTTACAACAGT |
| pBAD-VE-F1 | GGGCTAACAGGAGGAATTAACCATGAAGCGTTTGATGTTTATCGGCCCCA |
| pBAD-VE-R1 | CTTGCTCACGAATTCGCCAGAACCAGCAGCGGAGCCAGCGGATCCTTTTGTAAGACATAA |
| pBAD-K-F1 | GGGCTAACAGGAGGAATTAACCATGGCGAATAAGGAGCACCGCGT |
| pBAD-K-R1 | CTTGCTCACGAATTCGCCAGAACCAGCAGCGGAGCCAGCGGATCCCGCTTCACCTCGCTT |
| pBAD-M-F1 | GGGCTAACAGGAGGAATTAACCATGAACGGCGAAACCCTGCAGCGCA |
| pBAD-M-R1 | CTTGCTCACGAATTCGCCAGAACCAGCAGCGGAGCCAGCGGATCCCTCCTGCTTAATTAA |
| pBAD-N-F1 | GGGCTAACAGGAGGAATTAACCATGCATCTGGCACGAGTCACGGG |
| pBAD-N-R1 | CTTGCTCACGAATTCGCCAGAACCAGCAGCGGAGCCAGCGGATCCACACGAAAGCGTATC |

| | |
|---|---|
| pBAD-T-F1 | GGGCTAACAGGAGGAATTAACCATGTCTCAGGCTATAGGAATTTTAGA ACTCACCA |
| pBAD-T-R1 | CTTGCTCACGAATTCGCCAGAACCAGCAGCGGAGCCAGCGGATCCCCCC TCCACCATCTG |
| pBAD-U-F1 | GGGCTAACAGGAGGAATTAACCATGGAAAGACAACCGACAACGGATCG C |
| pBAD-U-R1 | CTTGCTCACGAATTCGCCAGAACCAGCAGCGGAGCCAGCGGATCCCGTC CGGGTGATCGA |
| pBAD-B-F1 | GGGCTAACAGGAGGAATTAACCATGAGCAGCAATGAGCTGGTGGAAC |
| pBAD-B-R1 | CTTGCTCACGAATTCGCCAGAACCAGCAGCGGAGCCAGCGGATCCTCA GATGTAGGACGG |
| pBAD-B'-F1 | GGGCTAACAGGAGGAATTAACCATGGCAGAAAAAAGCTGCAGTTTAAC GG |

**Table S3. The distribution of the number of "Shell first", "Cargo first", and "Concomitant" events in PduE-mCherry/ PduA-sfGFP and PduE-mCherry/ PduA-sfGFP assembly.** Note: The "Concomitant" event reported here either represents concomitant assembly, or a fusion following a shell/cargo independent assembly event.

| | **Shell first** | **Cargo first** | **Concomitant** |
|---|---|---|---|
| PduE-mCherry/PduA-sfGFP | 71 | 72 | 175 |
| PduJ-mCherry/PduE-sfGFP | 75 | 74 | 178 |

**Table S4. Diffusion coefficient and mobile fractions of Pdu MCP per cell measured using FRAP.** *n* represents the number of cells.

| | **Mobile fraction (%)** | **Diffusion coefficient ($D$, × $10^{-4}$ $\mu m^2 \cdot s^{-1}$)** | **Half Life ($\tau_{1/2}$, s)** |
|---|---|---|---|
| PduE (cargo) | 83 ± 6 ($n$ = 20) | 4.02 ± 1.89 ($n$ = 20) | 323 ± 171 ($n$ = 20) |
| PduA (shell) | 6 ± 3 ($n$ = 19) | 0.28 ± 0.15 ($n$ = 19) | 485 ± 407 ($n$ = 19) |
| PduGH (free) | 96 ± 4 ($n$ = 15) | N/A | 1.2 ± 0.4 ($n$ = 15) |
